# Cyclic-thiourea ionizable lipids program multilamellar mRNA-LNPs for sustained expression, frozen-storage stability and potent vaccine immunotherapy

**DOI:** 10.64898/2026.09.07.749752

**Authors:** Lei Zhang, Zhicheng Huang, Hao Gu, Ningze Zheng, Chunying Chen, Rong Cai, Aimin Hui

**Author notes:** These authors contributed equally: Lei Zhang, Zhicheng Huang, Hao Gu.

## Abstract

Lipid nanoparticles (LNPs) are the leading delivery vehicles for mRNA therapeutics, yet transient expression and limited formulation stability constrain their broader therapeutic application. Here we report the phenotype-driven discovery of an ionizable lipid (HI-62) bearing a 1,3,5-triazinane-2-thione (cyclic-thiourea) headgroup that reproducibly assemble mRNA into onion-like multilamellar nanoparticles. Cryogenic electron microscopy revealed ordered internal lamellae, while molecular-dynamics simulations supported a cooperative assembly model involving pH-responsive electrostatics, headgroup preorganization and short-range directional interactions. The lead HI-62 formulation maintained greater than 95% mRNA encapsulation across N/P ratios of 3-7, prolonged reporter expression for substantially longer durations than benchmark formulations following intravenous and intramuscular delivery, and retains physicochemical integrity and in vivo potency after repeated freeze-thaw cycles and 18 months of storage at −20 °C. In the tumor models, HI-62 elicits robust cellular responses and achieved significant inhibition of tumor growth. These findings establish cyclic-thiourea headgroup chemistry as a design principle for programming lipid-RNA organization into multilamellar architecture, and provide a design principle for next-generation LNPs platform with durable protein expression and superior stability. The HI-62-based formulation has been utilized in mRNA cancer vaccine program which will enter clinical development in months.

## Introduction

Messenger RNA (mRNA) therapeutics have become a programmable modality for inducing transient protein expression in vivo, enabling rapid vaccine development, protein replacement, genome editing and cancer immunotherapy^1-3^. Their clinical success has been made possible by lipid nanoparticles (LNPs), which protect labile RNA cargoes from degradation, promote cellular uptake and facilitate cytosolic delivery^2, 4^. Yet the broader therapeutic use of mRNA remains limited by inefficient intracellular release, short-lived protein expression, formulation instability and dose-limiting inflammatory or systemic toxicities. These limitations are particularly consequential for vaccines and immunotherapies, where the magnitude, duration and anatomical context of antigen expression shape immune priming and therapeutic efficacy^3-6^.

Ionizable lipids are the functional core of current mRNA-LNPs. Their pH-dependent protonation enables mRNA encapsulation during formulation, minimizes permanent cationic charge at physiological pH and contributes to endosomal membrane disruption after uptake. Most advances in ionizable lipid design have focused on tuning apparent pKa, biodegradability, linker chemistry and hydrophobic tail geometry^7^. These strategies have yielded clinically successful LNPs and increasingly potent preclinical formulations. However, the molecular principles by which ionizable lipid headgroups organize mRNA inside nanoparticles remain incompletely understood. In many formulations, lipid-mRNA association is still treated primarily as charge neutralization between protonated amines and the phosphate backbone^8^. This view explains RNA encapsulation, but it does not fully account for how subtle changes in headgroup chemistry can produce large differences in supramolecular architecture, intracellular release kinetics, long-term stability and immune activity.

Recent structural and mechanistic studies have begun to reveal that mRNA-LNPs are not uniform delivery vesicles but heterogeneous soft-matter assemblies with composition-dependent internal organization. Depending on lipid chemistry, buffer conditions and RNA cargo, LNPs may contain segregated mRNA domains, lipid-associated RNA states, bleb-like structures or ordered internal phases^9^. Such nanoscale architecture can influence mRNA stability, endosomal processing and biological output, yet they are usually emergent rather than deliberately programmed^4, 8^. This gap is important because intracellular delivery remains inefficient: after endocytosis, only a small fraction of internalized RNA reaches the cytosol, whereas much of the cargo is retained, recycled or degraded in end lysosomal compartments^10^. Thus, a central challenge in mRNA nanomedicine is no longer only to encapsulate RNA efficiently, but to control how RNA is packed, protected and released over time.

Formulation stability represents a second major barrier. mRNA-LNPs products are vulnerable to hydrolysis, lipid degradation, aggregation and freeze-thaw-induced structural stress. Existing stabilization strategies, including buffer optimization, cryoprotectants and lyophilization, can improve storage profiles but are typically applied after nanoparticle assembly^4, 11^. A more fundamental strategy would be to encode stability into the nanoparticle architecture itself. We reasoned that if specific ionizable lipid headgroups could impose ordered lipid-mRNA packing, and they might generate LNPs that simultaneously protect mRNA during storage and release it gradually after cellular uptake.

Thiourea motifs provide a chemical basis for such a design principle. In supramolecular chemistry and organ catalysis, thiourea groups are widely used as directional hydrogen-bond donors for binding electron-rich or anionic substrates^12^. Compared with purely electrostatic amine-phosphate interactions, thiourea-mediated recognition may offer additional directionality, cooperativity and geometric constraint. We therefore hypothesized that incorporating a cyclic thiourea motif into an ionizable lipid headgroup could convert mRNA complexation from a predominantly charge-driven process into a cooperative supramolecular interaction involving electrostatics, hydrogen bonding, stacking and ordered lamellar self-assembly.

Here we report a class of cyclic thiourea-containing ionizable lipids that program mRNA into onion-like multilamellar LNPs. Through phenotype-driven screening and structure-guided lipid engineering, we identified HI-62, a lead lipid that reproducibly assembles mRNA into multilamellar nanoparticles distinct from conventional benchmark LNPs. We propose that the cyclic thiourea motif functions as a supramolecular mRNA-binding unit rather than a passive structural linker. The ionizable amine mediates pH-responsive electrostatic interactions, while the thiourea N-H groups may provide directional hydrogen bonding with phosphate oxygens along the mRNA backbone. The thiocarbonyl group may enhance hydrogen-bond donation, and the cyclic scaffold may preorganize the headgroup geometry to favor ordered lipid-mRNA packing.

This molecular design produces a distinct structural and functional phenotype. Cryogenic electron microscopy and molecular dynamics simulations support a model in which HI-62 promotes ordered lipid-mRNA packing and onion-like multilamellar LNPs assembly. The resulting particles organize mRNA within layered lipid-RNA structures. Functionally, HI-62 LNPs prolong protein expression after both systemic and intramuscular administration, preserve particle integrity and mRNA encapsulation after repeated freeze-thaw stress, and retain multi-lamellae and in vivo potency after storage at −20 °C for 18 months. In tumor vaccine models, this architecture translates into potent immune activation and robust therapeutic efficacy, including regression of established tumors. These findings identify cyclic thiourea as an ionizable lipid design motif for programming lipid-mRNA supramolecular interactions and establish onion-like multilamellar assembly as a controllable architectural parameter for durable mRNA nanomedicine.

## Results and discussion

### Formulation optimization and structural characterization of HI-62 LNPs

Phenotype-driven screening and structure-guided lipid engineering identified HI-62 as cyclic thiourea-containing ionizable lipid to reproducibly assembles mRNA into multilamellar nanoparticles (Fig. 1A). We optimized the physicochemical properties and transfection activity of HI-62 LNPs using a central-composite design coupled to response-surface modelling. The analysis nominated formulation E21 for subsequent studies. Microfluidic assembly produced particles with a hydrodynamic diameter of 89.3 ± 0.6 nm, low polydispersity of 0.03, a near-neutral zeta potential of 1.95 mV and an mRNA encapsulation efficiency of 97.5% (Fig. 1B). A 2- [ (4-methylphenyl) amino] naphthalene-6-sulfonic acid (TNS) assay yielded an apparent particle pKa of 6.45 (Fig. 1B). This value is within the range often associated with productive mRNA delivery, but pKa alone does not establish the efficiency or route of endosomal escape^13, 14^.

**Figure 1.**
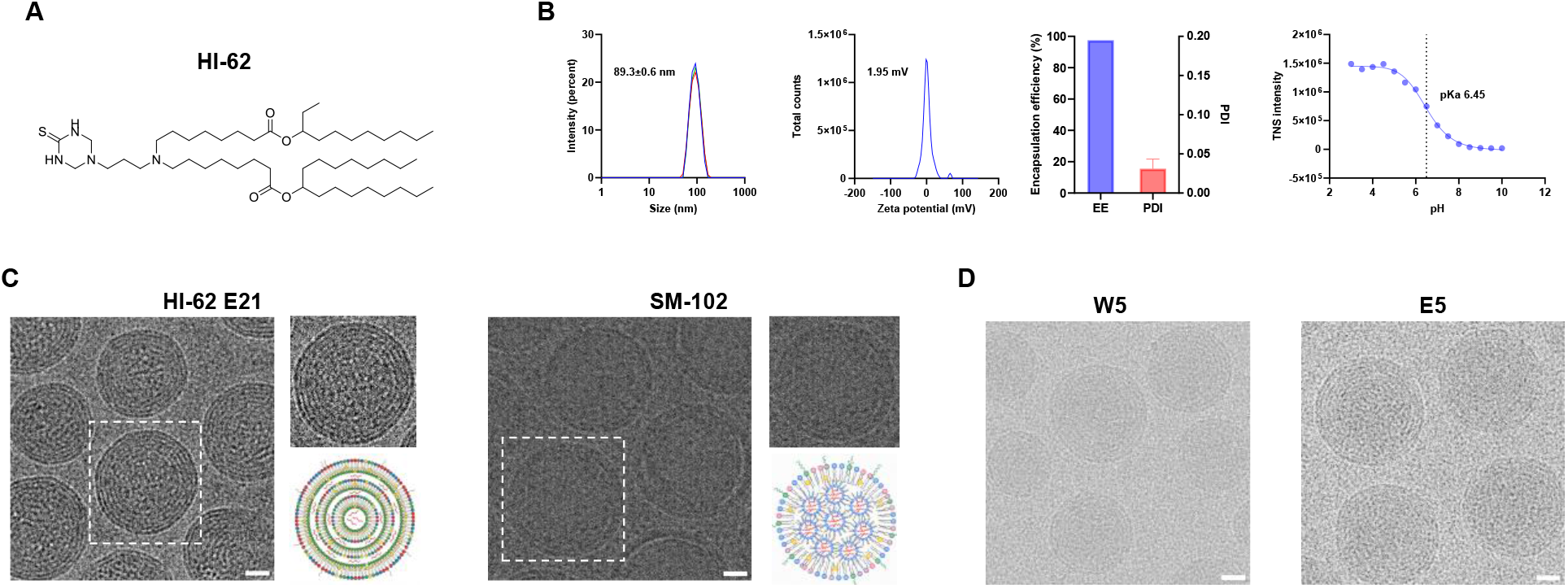
Phenotype-driven discovery of cyclic-thiourea ionizable lipids that program multilamellar mRNA-LNPs assembly. (A) Chemical structures of HI-62 and the SM-102 comparator. (B) Physicochemical characterization of E21 LNP formulation, including particle-size distribution, zeta potential, polydispersity, encapsulation efficiency and TNS-derived apparent pKa. (C) Representative cryo-EM images comparing the electron-dense internal morphology of SM-102 LNPs with the concentric lamellae of HI-62 LNPs, scale bars are 20 nm. (D) Representative cryo-EM images of HI-62 across the indicated formulation variants.

Cryo-EM revealed a multilamellar, onion-like internal morphology for HI-62 LNPs, in contrast to the predominantly electron-dense amorphous cores observed for the SM-102 benchmark (Fig. 1C). During acidic microfluidic mixing, protonation of the HI-62 amine can support electrostatic association with the mRNA phosphate backbone. The cyclic-thiourea headgroup additionally presents donor and acceptor groups in a constrained geometry, providing a plausible basis for short-range directional interactions. Across the tested helper-lipid species and molar-ratio variants, i.e. W5 and E5, HI-62-containing particles retained the multilamellar phenotype (Fig. 1D), supporting HI-62 as a principal determinant of internal organization while not excluding contributions from the other formulation components. The structural data and simulations below are interpreted as convergent support for a cooperative assembly model, rather than as direct proof of persistent hydrogen bonds or stacking interactions in intact particles.

### Multilamellar organization is associated with efficient encapsulation and prolonged expression

HI-62 LNPs maintained mRNA encapsulation efficiencies above 95% across N/P ratios of 3-7 and achieved 97% encapsulation at N/P 3 (Fig. 2A). These data show efficient loading across a broad electrostatic stoichiometry. The multilamellar architecture may contribute to this efficient loading by providing repeated lipid-aqueous interfaces that could accommodate and organize mRNA more effectively^15^. However, direct measurements of intraparticle RNA distribution and aqueous-volume fraction would be required to establish this structural interpretation.

**Figure 2.**
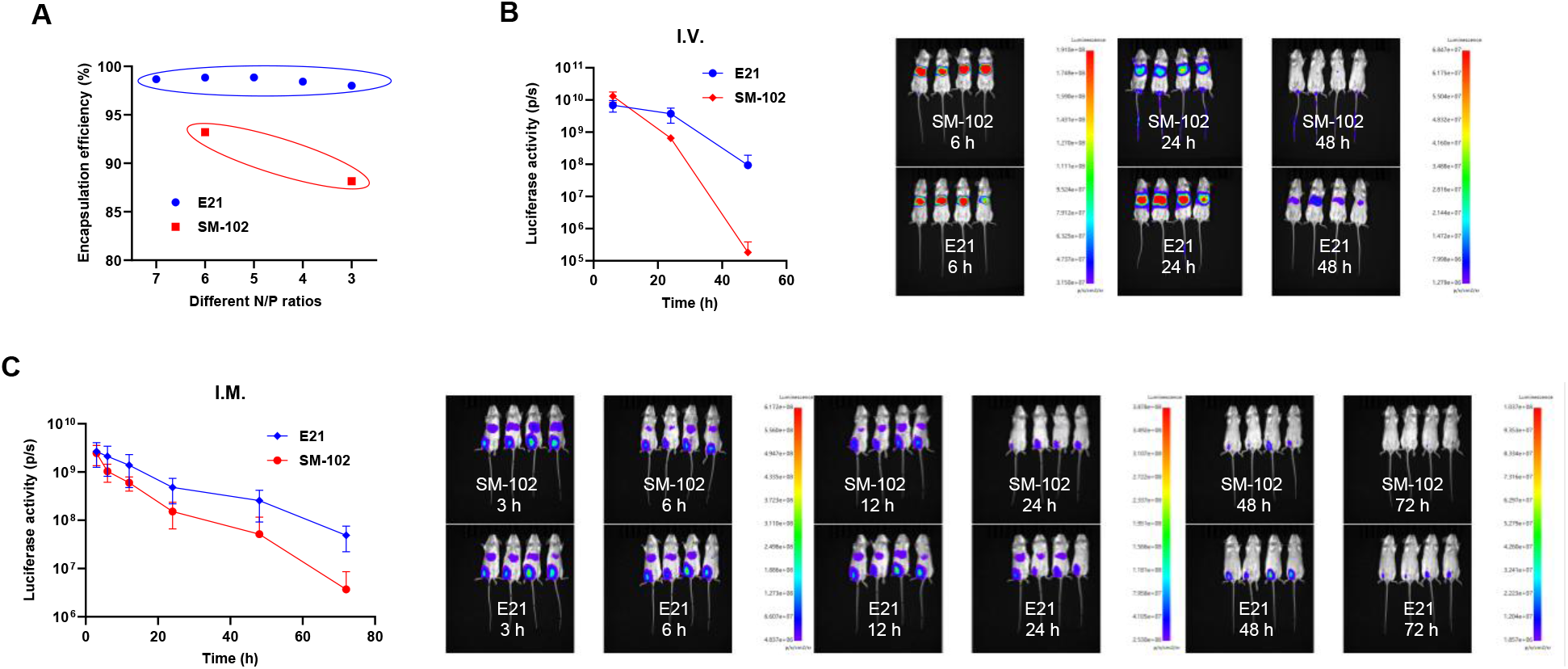
Multilamellar HI-62 LNPs efficiently encapsulate mRNA and prolong reporter expression in vivo. (A) mRNA encapsulation efficiency of HI-62 and SM-102^18^ formulations across the indicated N/P ratios. (B) Longitudinal whole-animal reporter activity after intravenous administration, with representative images at the indicated times. (C) Reporter-expression kinetics and representative images after intramuscular administration.

We next asked whether the structural difference was accompanied by expression kinetics in vivo. HI-62 and SM-102 LNPs were administered intravenously and bioluminescence was monitored. Reporter signals were comparable at 3 h after injection, but by 48 h the SM-102 signal approached baseline whereas the HI-62 signal remained more than three orders of magnitude higher (Fig. 2B). Intramuscular administration showed a similarly extended expression profile (Fig. 2C). These route-consistent data associate the HI-62 formulation and its multilamellar phenotype with sustained translation. One possible explanation is that the repeated interfaces in multilamellar particles may enable more gradual membrane disruption and mRNA release following endosomal uptake.^16, 17^ Direct measurements of endosomal membrane disruption and intracellular mRNA release are needed to test this proposed mechanism.

### HI-62 LNPs retain physicochemical attributes and potency during frozen storage

We evaluated whether the distinctive HI-62 organization was accompanied by resistance to freeze-thaw stress. After six freeze-thaw cycles, the encapsulation efficiency of SM-102 LNPs decreased from 96% to below 90%, whereas HI-62 LNPs remained above 95%. Meanwhile, the hydrodynamic size of SM-102 formulation showed slight increase under freeze-thaw stress (Fig. 3A). These results demonstrate retention of encapsulation and size of HI-62 under the tested conditions.

**Figure 3.**
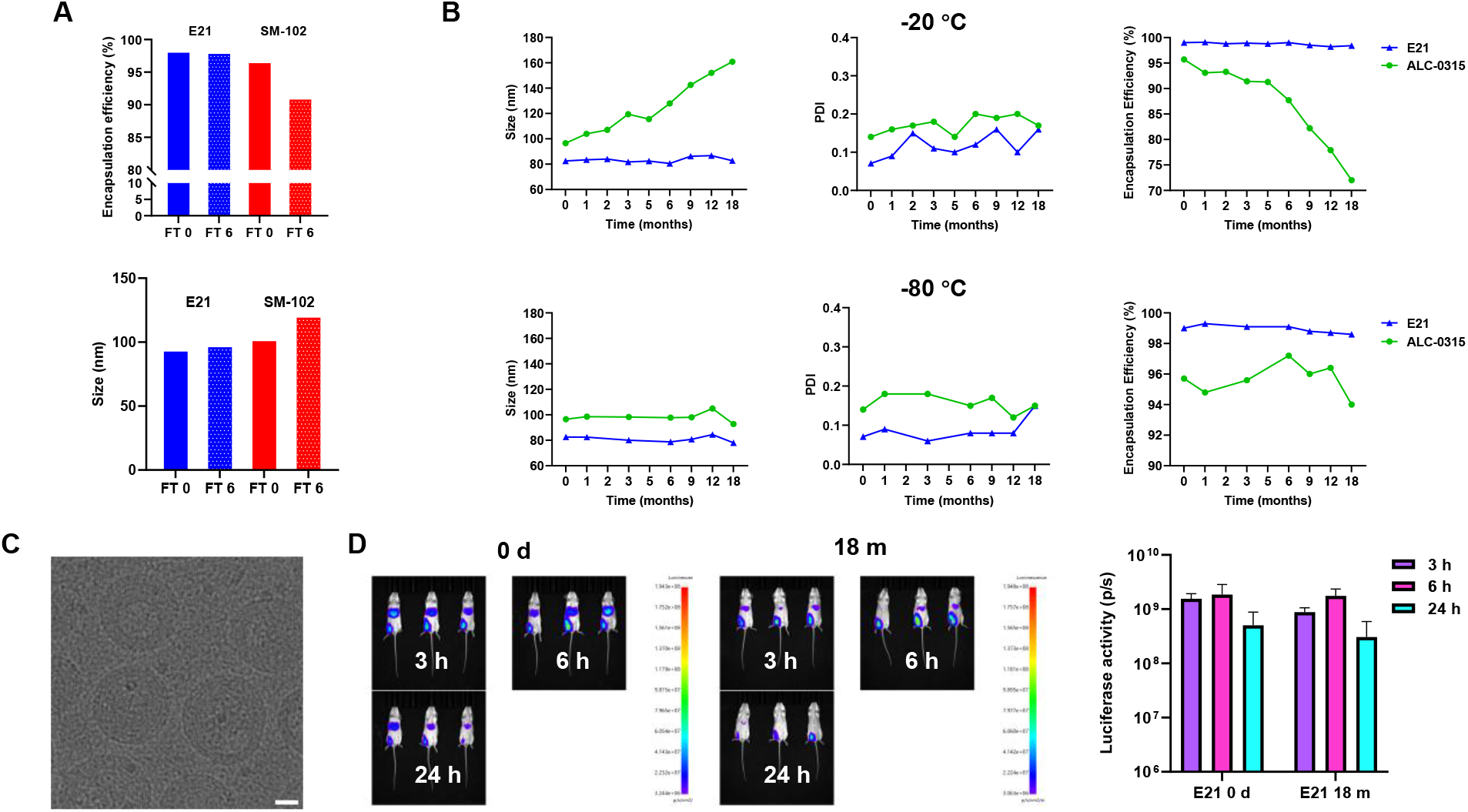
HI-62 LNPs retain particle attributes, internal order and biological potency during frozen storage. (A) mRNA encapsulation efficiency and hydrodynamic size before and after six freeze-thaw cycles for HI-62 and indicated benchmark formulation. (B) Longitudinal particle size, PDI and encapsulation efficiency at −20 °C and −80 °C for HI-62 and the indicated benchmark formulation. (C) Representative cryo-EM image after frozen storage at −20 °C for 18 months, scale bar is 20 nm. (D) Representative in vivo reporter images comparing stored and fresh formulations.

We then compared real-time frozen-storage profiles of HI-62 and ALC-0315 LNPs. At −20 °C, the mean HI-62 particle diameter changed from approximately 80 to 83 nm, whereas the ALC-0315 comparator increased to approximately 160 nm by the final reported time point. Encapsulation remained at 97% for HI-62 and decreased to 77% for ALC-0315; both formulations retained their measured particle attributes at −80 °C (Fig. 3B). The size increase and loss of encapsulation in the comparator are consistent with colloidal and cargo-retention instability.

Cryo-EM of HI-62 LNPs recovered after 18 months storage at −20 °C showed retention of concentric lamellae (Fig. 3C). Stored formulations produced in vivo reporter activity comparable to freshly prepared material under the tested conditions (Fig. 3D). Together, these orthogonal measurements support HI-62 LNPs preserve structure and biological potency during frozen storage. The multilamellar architecture may contribute to this stability by providing multiple lipid layers that can accommodate structural perturbations during freeze-thaw cycling,^19^ although the underlying mechanism remains to be elucidated.

### Molecular simulations support a cooperative model of multilamellar assembly

To understand why HI-62 reproducibly assembles mRNA into multilamellar structures while conventional lipids do not, we performed molecular docking^20^ and molecular-dynamics (MD) simulations of HI-62 and SM-102 interactions with RNA-containing assemblies. Molecular docking revealed a hydrogen bond between the HI-62 cyclic-thiourea headgroup and the RNA backbone with docking score of −2.452 kcal/mol, whereas no hydrogen bonding was detected between SM-102 and RNA. (Fig. 4A). In 10-ns MD simulations. HI-62 established a transient cyclic thiourea-base close contact at 1.040 ns, which may indicate a possible stacking interaction (Fig. 4B).

**Figure 4.**
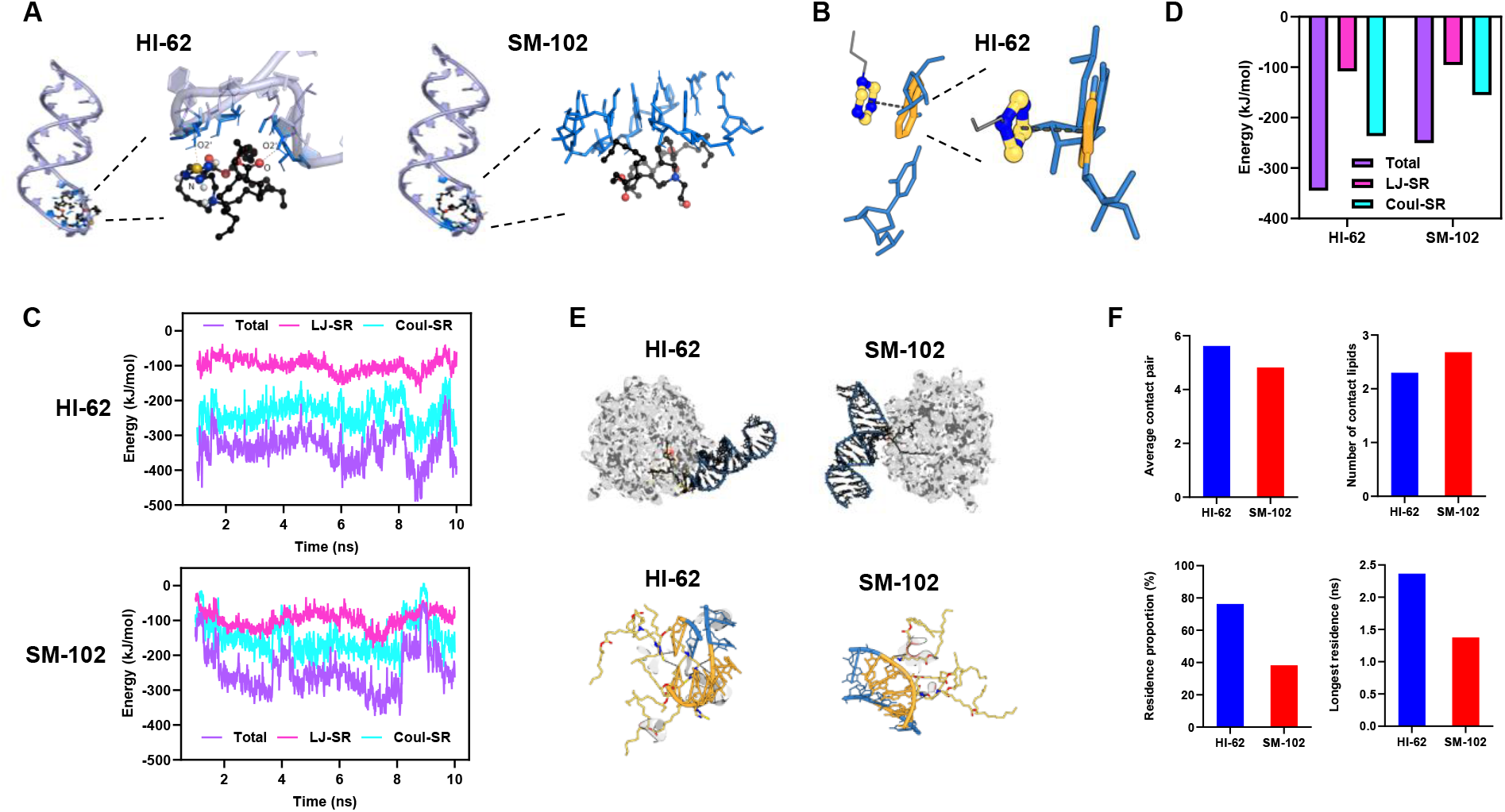
Molecular docking and molecular-dynamics support cooperative cyclic-thiourea-mediated lipid-RNA organization. (A) Representative docking configurations for HI-62 and SM-102 with RNA, highlighting candidate headgroup-phosphate. (B) Representative transient close contact of headgroup-base stacking. (C) Profiles of RNA-lipid short-range nonbonded interaction energies over 1-10 ns, including total, van der Waals, and Coulombic components. (D) Quantitative analysis of average interaction energies over 5-10 ns. (E) Representative surface-contact conformation with RNA at 1.040 ns, with a magnified view of the local contact interface. RNA backbone is shown in blue, contacting nucleotides in orange, and lipids in element-based colors. (F) Quantitative comparison of the average number of phosphate-head-group contact pairs, average number of contacting lipids, RNA surface residence fraction, and longest continuous surface residence time.

We further analyzed the time-dependent profiles of short-range nonbonded interaction energies throughout the simulation^21^ and calculated their average values over the 5-10 ns (Fig. 4C, D). Coulombic interactions (Coul-SR) and Lennard-Jones interactions (LJ-SR) respectively represent electrostatic and van der Waals contributions. HI-62 exhibited substantially more negative interaction energy than SM-102, with 5-10 ns average values of −344.7 and −250.4 kJ/mol respectively, indicating stronger interactions between HI-62 and RNA. Headgroup contact profiles were next compared between HI-62 and SM-102. Representative lipid-RNA contact conformations at 1.040 ns showed a larger contact area for HI-62, with a greater proportion of the RNA backbone involved in the interface (Fig. 4E). Quantitative analyses of contact pair, lipid count, RNA residence further demonstrated more extensive and sustained HI-62-RNA interactions (Fig. 4F). Together, these findings support stronger HI-62-RNA interactions and provide convergence for the proposed cooperative assembly model.

Coarse-grained molecular dynamics (CG-MD) simulations were conducted in a local system containing an RNA layer between two lipid bilayers.^22-24^ HI-62 (+1) and HI-62 (+2) represent different protonation states. Representative snapshots at 0, 200, 800, and 1000 ns showed that multilayer organization maintained throughout the simulation (Fig. 5A). Consistent with these observations, RNA maintained extensive contacts with both the lower and upper inner leaflets, with average contact residues of 443.2 and 436.7 (Fig. 5B).

**Figure 5.**
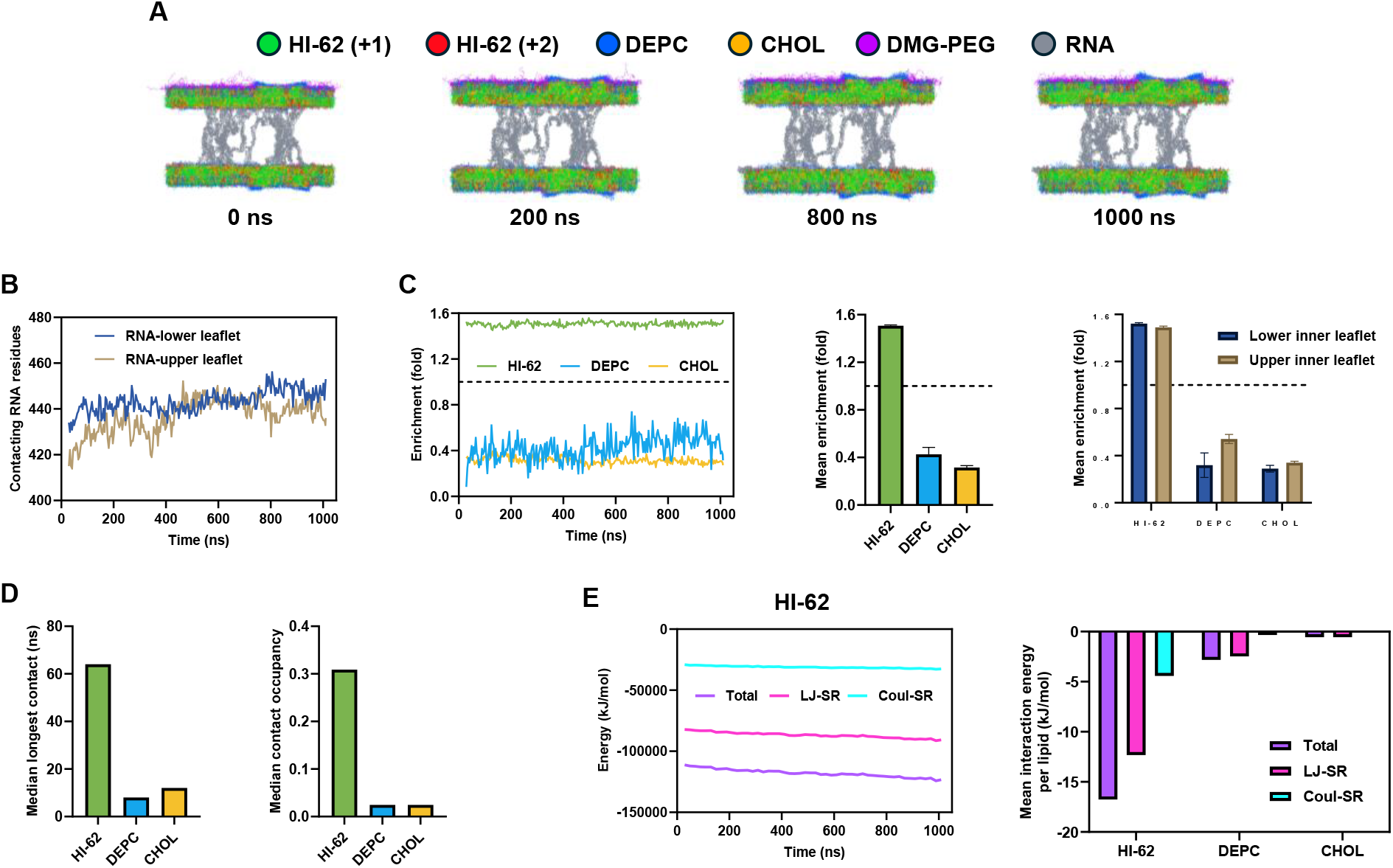
Coarse-grained molecular dynamics support preferential HI-62-RNA association at the membrane interface. (A) Representative CG-MD snapshots at 0, 200, 800, and 1000 ns showing the localization of RNA between the two inner membrane leaflets and the maintained multilayer organization. HI-62 (+1) and HI-62 (+2) represent different protonation states. (B) Quantitative analysis of RNA contacts with the lower and upper inner membrane leaflets. (C) Quantitative comparison of interfacial enrichment factors for HI-62, DEPC, and CHOL, including enrichment at the lower and upper interfaces and across consecutive time windows. (D) Quantitative comparison of the median longest continuous residence time and contact occupancy of HI-62, DEPC, and CHOL at the RNA interface. E, Quantitative comparison of RNA-lipid short-range nonbonded interaction energies.

We next analyzed lipid enrichment and residence behavior at the RNA interface. HI-62 showed the highest interfacial enrichment, with an average enrichment factor of 1.507, compared with 0.426 for DEPC and 0.315 for cholesterol (CHOL), and exhibited comparable enrichment at the lower and upper interfaces (1.522 and 1.490, respectively) (Fig. 5C). HI-62 also showed substantially longer and more frequent residence at the RNA interface, with a median longest continuous residence time of 64.0 ns and contact occupancy of 30.9%, compared with 8.0 and 12.0 ns and 2.4% and 2.4% for DEPC and CHOL, respectively (Fig. 5D). RNA-lipid nonbonded interaction energies further showed that HI-62 had the strongest interaction with RNA, average per-lipid total interaction energies of −16.751, −2.817, and −0.556 kJ/mol for HI-62, DEPC, and CHOL (Fig. 5E). Because trajectory stability is not a direct thermodynamic measurement and the model represents a restricted system, these simulations are used to generate and support a mechanistic hypothesis rather than to establish causality. Collectively, the computational and structural results support a model in which electrostatics, cyclic-headgroup geometry and short-range interactions favor multilamellar organization.

### A HI-62-based therapeutic vaccine controls established tumors

We next evaluated the antitumor activity of the HI-62-based mRNA-LNPs formulation in mice bearing established tumors. After tumor establishment, mice received three doses of HI-62 or ALC-0315 LNPs at 5-day intervals, followed by monitoring of tumor growth and assessment of antigen-specific IFN-γ responses (Fig. 6A). Compared with the Tris control group, both HI-62- and ALC-0315-based formulations significantly suppressed tumor growth, with the HI-62 LNPs achieving a higher proportion of complete tumor responses (4 of 5 mice) (Fig. 6B). Meanwhile, HI-62 induced a higher frequency of antigen-specific IFN-γ producing CD8^+^ T cells in peripheral blood than ALC-0315, although with no statistical significance (Fig. 6C). Together, these findings demonstrate that the HI-62-based LNPs formulation elicits pronounced antitumor activity accompanied by robust antigen-specific cellular immune responses.

**Figure 6.**
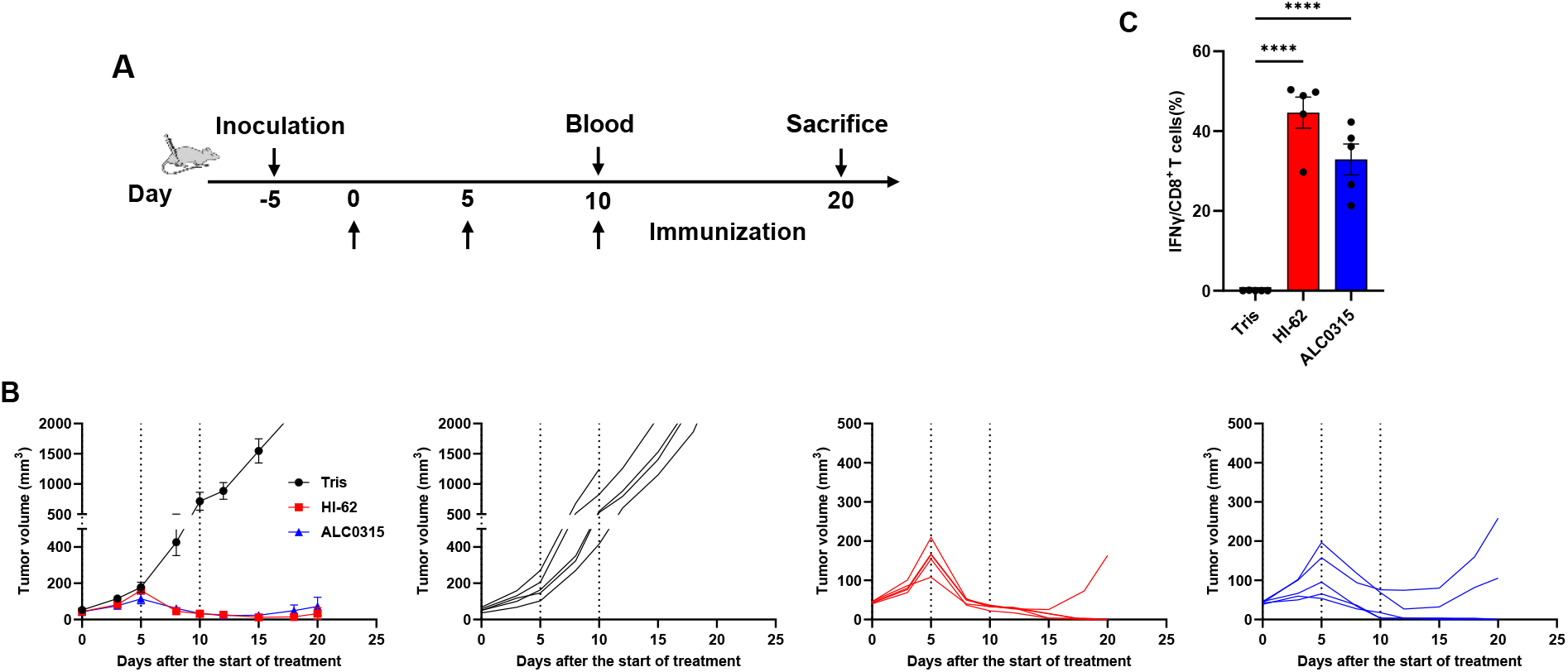
HI-62-based therapeutic vaccine formulation controls established TC-1 tumors and elicits antigen-specific cellular responses. (A) Therapeutic vaccination schedule in mice bearing established tumors. Mice were enrolled and received three doses of mRNA-LNPs (10 μg mRNA per dose) via intramuscular injection on days 0, 5, and 10 after tumor inoculation. Groups included Tris vehicle control, HI-62 LNPs, and ALC-0315 LNPs (n = 5). Peripheral blood were collected on days 10, and all mice were euthanized on day 20 unless the tumor volume reached the ethical endpoint criteria earlier. (B) Mean and individual tumor-volume trajectories for the Tris, HI-62, and ALC-0315 groups; dashed lines indicate treatment times. Mice were sacrificed upon reaching a tumor volume of 2000 mm^3^. (C) Frequency of antigen-specific IFN-γ^+^ CD8^+^ T cells in peripheral blood. Blood cells were isolated, stimulated with specific peptide, surface and intracellularly stained and analyzed by flow cytometry. Statistical significance was assessed using One-way ANOVA. ****P < 0.0001.

### Exploratory secondary-lymphoid-organ targeting

Given that efficient delivery to secondary lymphoid organs,namely, the spleen and lymph nodes, is critical for the therapeutic efficacy of mRNA-based interventions in immune-related diseases,^25-27^ we investigated whether modulating HI-62 formulation parameters could redirect systemic mRNA expression toward these tissues. Compared with the liver-predominant E21 formulation, variants P19 and P20 markedly shifted systemic mRNA expression toward the spleen, which accounted for 72% and 60% of total measured expression, respectively, resulting in an approximately sixfold increase in splenic bioluminescence relative to E21 (Fig. 7A). P19 and P20 also elicited significantly higher frequencies of IFN-γ^+^ CD8^+^ T cells than E21 in both blood and spleen (Fig. 7B), supporting their enhanced capacity to induce antigen-specific cellular responses. We next sought to further enhance splenic targeting by incorporating anionic lipids into the P20 formulation.^28, 29^ Addition of 2 or 4 mol% phosphatidic acid (PA), phosphatidylglycerol (PG), or phosphatidylserine (PS) further modulated the organ distribution of mRNA expression. Among these variants, P20 + 4PS produced the highest splenic luminescence, corresponding to an approximately sevenfold increase relative to P20 alone, and increased the splenic fraction of total measured expression to more than 90% (Fig. 7C). Notably, P20 and all anionic lipid-containing variants also generated detectable expression in the lymph nodes, with the PS-containing formulations producing particularly strong lymph-node signals (Fig. 7C). Together, these iterative formulation studies demonstrate that HI-62 provides a tunable platform for redirecting systemic mRNA expression toward secondary lymphoid organs.

**Figure 7.**
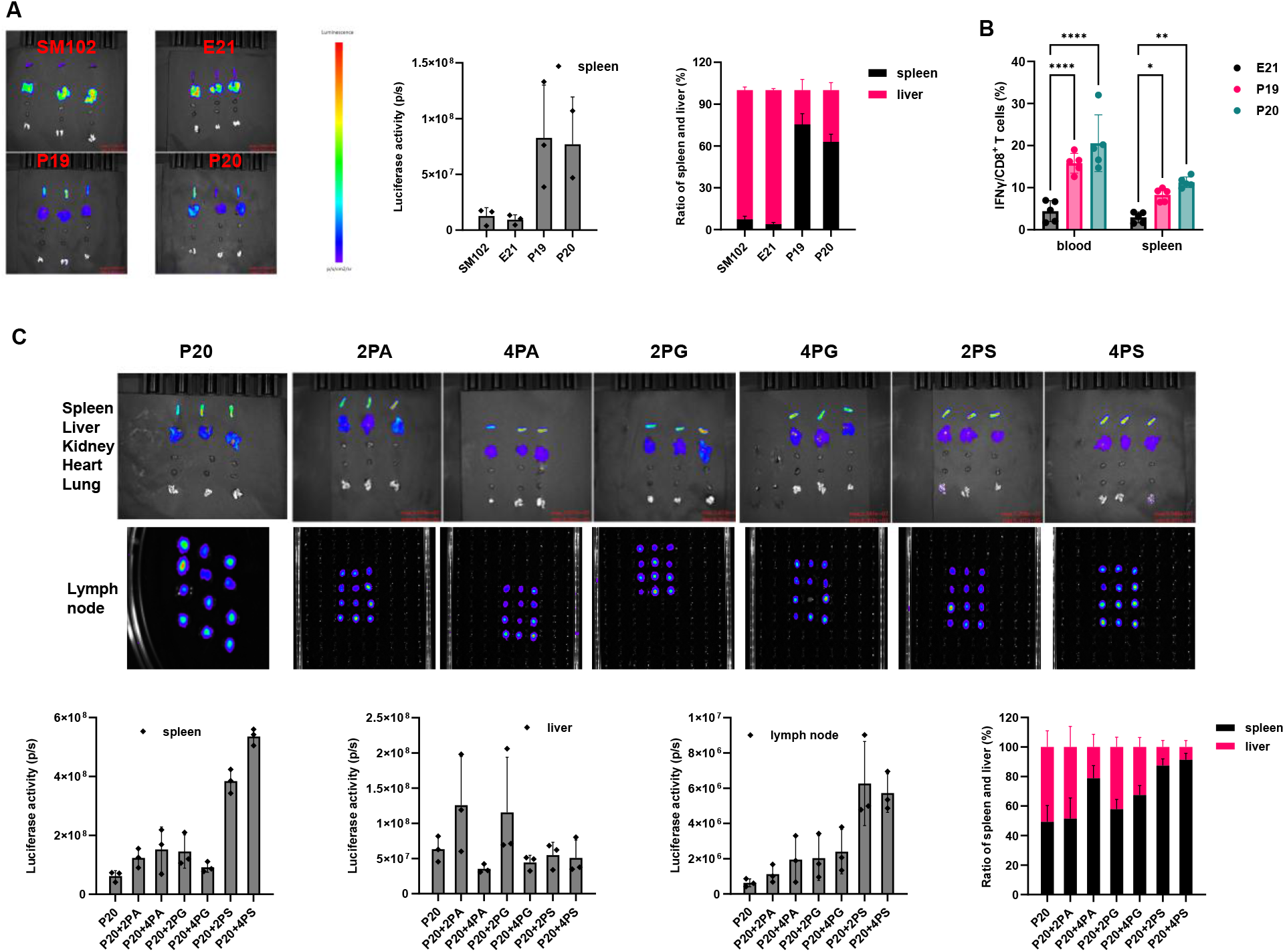
Exploratory organ targeting of HI-62-based mRNA formulations. (A) Ex vivo bioluminescence imaging of organs following administration of the indicated mRNA formulations (5 μg per mouse). D-luciferin potassium salt (150 mg kg^−1^) was administered 6 h after mRNA administration, and organs were collected and imaged 10 min later. Quantification of total splenic bioluminescence and the relative distribution of bioluminescence between the spleen and liver is shown (n = 3). (B) Frequencies of IFN-γ^+^ CD8^+^ T cells in blood and spleen following administration of the indicated formulations. Mice received 10 μg of mRNA on days 0, 5 and 10 (n = 5). Peripheral blood was collected on day 15, followed by spleen collection on day 16 for analysis. (C) Ex vivo bioluminescence imaging of organs following incorporation of the indicated anionic lipids into P20 was conducted as described above. Quantification of luciferase activity in the spleen, liver and lymph nodes was analyzed. The relative distribution of bioluminescence between the spleen and liver is also calculated (n = 3). Statistical significance was determined by two-way ANOVA. *P < 0.05, **P < 0.01 and ****P < 0.0001.

## Conclusion

Here we identify cyclic-thiourea headgroup chemistry as a strategy for programming the supramolecular organization of mRNA lipid nanoparticles. The lead lipid, HI-62, reproducibly formed onion-like multilamellar nanoparticles with efficient mRNA encapsulation, prolonged protein expression and retained activity after extended frozen storage. Computational analyses support a cooperative assembly model involving pH-responsive electrostatics, headgroup preorganization and short-range interactions. HI-62-based formulations elicited potent immune responses and marked suppression of established tumors. Formulation optimization further redirected systemic mRNA expression towards secondary lymphoid organs. Together, these findings establish a next generation mRNA-LNPs platform with engineered durability, tissue distribution and therapeutic performance. The HI-62-based formulation has been utilized in mRNA cancer vaccine program which will enter clinical development in months.

## Methods

### Study design

This study investigated the relationship between ionizable-lipid headgroup chemistry, internal organization, physicochemical stability and biological performance of mRNA lipid nanoparticles (LNPs). The study comprised four main stages: identification and formulation optimization of cyclic-thiourea-containing ionizable lipids; physicochemical and structural characterization of the lead HI-62 formulation; evaluation of reporter expression and frozen-storage stability; and assessment of HI-62-based formulations in tumor suppression and secondary-lymphoid-organ targeting models.

### mRNA preparation

Firefly luciferase (FLuc) mRNA, antigen mRNA were prepared by in vitro transcription from the corresponding linearized DNA templates. The mRNA transcripts were generated using the specified untranslated-region and poly(A) designs and incorporated modified nucleosides as indicated for each construct. Transcripts were capped using the corresponding capping procedure and purified before formulation. Unless otherwise stated, the same mRNA preparation was used for head-to-head comparisons within an individual experiment.

### LNPs formulation

HI-62, helper phospholipid, cholesterol and DMG-PEG2000 were dissolved in ethanol at the indicated molar ratios. mRNA was diluted in citrate buffer. The lipid-containing organic phase and mRNA-containing aqueous phase were rapidly mixed using a microfluidic mixing device under 1:3 organic to aqueous volume ratio at a total flow rate of 20 mL/min. Following mixing, the resulting LNPs suspension was diluted and buffer exchanged to remove ethanol and adjust the formulation to the final buffer conditions. Unless otherwise stated, LNPs were subsequently concentrated and stored at the indicated temperature before characterization or administration. N/P was defined as the molar ratio of HI-62 molecule to mRNA phosphate groups.

### Particle size, polydispersity, zeta potential, apparent pKa and encapsulation efficiency

Hydrodynamic diameter and polydispersity index (PDI) were measured by dynamic light scattering (DLS); zeta potential was measured by electrophoretic light-scattering (ELS), both using Zetasizer Lab (Malvern Panalytical).

The apparent pKa of LNPs was determined using a TNS fluorescence assay. LNP samples were incubated with TNS over pH 3.0 to 10.0, and fluorescence intensity was measured under 321/445 nm. Normalized fluorescence values were fitted to a sigmoidal response model, and the midpoint of the fitted curve was reported as the apparent particle pKa.

mRNA encapsulation efficiency was determined using a RiboGreen-based fluorescence assay (Thermo Fisher). Encapsulation efficiency was calculated as: Encapsulation efficiency (%) = 100 × (total RNA − unencapsulated RNA) / total RNA.

### Cryo-EM and morphology analysis

LNP samples were prepared for cryogenic transmission electron microscopy (cryo-EM) using standard vitrification procedures. Briefly, an aliquot of LNP suspension was applied to EM grid, blotted under controlled temperature and humidity conditions and rapidly vitrified in liquid ethane. Grids were examined using a cryo-electron microscope operated under low-dose conditions. Representative images were acquired using identical or appropriately matched imaging conditions for comparative experiments.

The E21 and W5/E5 formulations were imaged using different cryo-electron microscopy systems, resulting in differences in image brightness and contrast. These differences are attributable to the imaging systems and do not affect the assessment of particle morphology.

### In vivo reporter expression

Female BALB/c mice were used for in vivo reporter-expression studies. Mice received 5 μg of FLuc mRNA per animal formulated in the indicated LNPs by intravenous or intramuscular administration. At the indicated time points after administration, mice were injected with D-luciferin potassium salt and imaged using an in vivo bioluminescence imaging system (AniView100, Biolight Biotechnology). Imaging parameters and exposure settings were kept consistent within each experiment. Bioluminescence was quantified as total flux from predefined anatomical regions of interest.

### Freeze-thaw stability and frozen-storage studies

HI-62 and SM-102 LNPs were subjected to six repeated freeze-thaw cycles under the experimental conditions. Before and after the treatment, particle size and mRNA encapsulation efficiency were determined using the methods described above.

For long-term storage studies, LNP formulations were stored at −20 °C or −80 °C. Samples were analyzed at the indicated time points for particle size, PDI and mRNA encapsulation efficiency. For the extended −20 °C storage study, HI-62 LNPs were evaluated after 18 months of storage. Cryo-EM was used to examine the characteristic multilamellar morphology. Stored and freshly prepared formulations were subsequently compared in vivo for reporter expression.

### Molecular docking and MD simulation

Molecular docking was performed using AutoDock Vina 1.2.0 with a local RNA structure as the receptor and the pH 4 structures of Lipid as ligands.^20^ The RNA-lipid systems were docked under identical conditions, and the lowest-scoring pose for each lipid was selected for structural analysis. RNA-lipid interactions were identified using PLIP. Molecular structures were visualized using PyMOL.

All-atom molecular dynamics simulations were performed using a 37-nt RNA with a surface model containing lipid/DOPC/CHOL. RNA, DOPC/CHL and lipid were parameterized using the Amber OL3, Amber Lipid21 and GAFF2 force fields, respectively.^21^ The systems were solvated with TIP3P water, neutralized and supplemented with approximately 0.15 M NaCl. Simulations were performed at 300 K and 1 bar using V-rescale temperature coupling and isotropic Parrinello-Rahman pressure coupling, with a 2-fs integration step. Long-range electrostatics were treated using particle-mesh Ewald, with 1.0-nm cutoffs for short-range electrostatic and van der Waals interactions and long-range dispersion corrections.

Coarse-grained molecular dynamics simulations were performed using Martini 3.0.0 in GROMACS 2021.5.^21, 30^ The system comprised a 1,973-nt RNA positioned between two HI-62/DEPC/CHOL bilayers (∼40 × 40 nm), with HI-62 accounting for certain proportion of membrane lipids. The system was solvated with Martini water and neutralized with counterions. After energy minimization and equilibration, 1-μs production simulations were performed at 300 K using a 0.020-ps timestep and V-rescale thermostat. Electrostatic and van der Waals interactions used 1.1-nm cutoffs. Trajectories were analysed using GROMACS and Python and visualized with PyMOL.

### Anti-tumor study and cellular analyses

C57BL/6 mice with established tumors received three intramuscular administrations of mRNA-LNPs at 5-day intervals, with 10 μg mRNA per dose. Tumor dimensions were monitored at the indicated intervals and tumor volume was calculated. Peripheral blood was collected on day 10 to assess antigen-specific cellular immunogenicity, and antigen-specific IFN-γ-producing CD8^+^ T cells were quantified by flow cytometry following stimulation with the relevant antigen.

### Secondary-lymphoid-organ targeting

Organ targeting. SM-102, E21, P19 and P20 LNPs encapsulating reporter mRNA were administered intravenously at 5 μg mRNA per mouse. Six hours after administration, D-luciferin potassium salt (150 mg kg^−1^) was administered, followed by organ collection and ex vivo bioluminescence imaging 10 min later. Bioluminescence was quantified in the spleen and liver. Cellular immune response. Mice received 10 μg mRNA on days 0, 5 and 10 by the indicated route. Peripheral blood was collected on day 15 and spleens on day 16. Antigen-specific IFN-γ^+^ CD8^+^ T cell frequencies were determined by flow cytometry following antigen stimulation.

Anionic-lipid modification. P20 formulations were supplemented with 2 or 4 mol% phosphatidic acid (PA), phosphatidylglycerol (PG) or phosphatidylserine (PS) and administered intravenously. Ex vivo bioluminescence was measured in the spleen, liver and lymph nodes using the same imaging procedure.

## Statistical analysis

Data are presented as mean ± standard deviation (s.d.) based on independent experiments. Statistical significance was determined using GraphPad Prism and stated in each figure legend.

## Data availability

All data supporting the plots are available within the paper, or from the authors upon reasonable request.

## Acknowledgements

This work was supported by Special Fund for the China-Israel Biotech Industry Incubation Base of Guangzhou Development District, National Key R&D Program of China (2024YFC2310500),New Cornerstone Science Foundation, National Natural Science Foundation of China (22595380), Basic Science Center Project of the National Natural Science Foundation of China (22388101).

## Author contributions

L.Z., Z.H., and H.G. conceived the work, performed the experiments, analyzed the data and wrote the paper. N.Z. contributed to the anti-tumor experiments. C.C. contributed to manuscript writing and revision. R.C. and A.H. conceived and supervised the project and revised the paper. The final paper was approved by all authors.

